# Suppression of electron transport chain Complex III assembly is an adaptive mechanism to reduce host inflammation during viral infection

**DOI:** 10.64898/2026.01.30.702752

**Authors:** Sheryl Beh, Cheryl Q.E. Lee, Kimberle Shen, Chengwei Zhong, Chinmay K. Mantri, Liang Chao, Radiance Lim, Tan Kiat Yi, Radoslaw M. Sobota, Jinmiao Chen, Ashley L. St John, Lena Ho

## Abstract

Mitochondrial electron transport and oxidative respiration are required for immunity and host tolerance. How the ETC contributes to viral infections - where a proinflammatory anti-viral response needs to be finely balanced with host-protective anti-inflammatory mechanisms - remains unclear. Here, we demonstrate that following infection by H1N1 influenza virus, murine macrophages reduce Complex III levels by downregulating the early CIII assembly factor called the COMB complex. To determine the effect of CIII suppression, we utilized Brawnin (Br or UQCC6) knockout mice, which lack COMB complexes and have reduced Complex III. Following H1N1 infection, Br KO bone marrow-derived macrophages (BMDMs) had reduced inflammation due to reduced CIII Qo site ROS. Br KO mice infected with a non-lethal dose of H1N1 had reduced lung immune pathology, viral burden and enhanced recovery following H1N1 infection. Single-cell RNAseq analysis revealed that H1N1-infected Br KO lungs had reduced abundance of hyper-inflammatory monocytes known to cause severe respiratory disease and reduced monocyte chemotaxis. Our study demonstrates that ETC Complex III suppression is part of an innate immune response to viral infection, and is a potential strategy to control host inflammation in acute respiratory viral infections.

## Introduction

Mitochondria, traditionally recognized as the cellular powerhouses involved in energy production through oxidative phosphorylation, have emerged as integral players of immune cell fate and function, highlighting the importance of cellular bioenergetics in shaping the immune landscape. The electron transport chain (ETC), residing within the inner mitochondrial membrane, comprises of 4 complexes – Complex I, II, III, and IV that couple electron transfer from fuel to oxygen with proton extrusion to build a proton motive force that drives ATP synthesis at Complex V. Beyond its canonical role in ATP production, the ETC exerts regulatory influence on both adaptive and innate immunity. In the adaptive branch, T helper cell development and effector functions are controlled by the interplay between Complex I and II. While Complex I and aspartate production drive proliferation of T helper cells, Complex II activity antagonizes differentiation and enforces terminal effector function (Bailis et al., 2019). Following activation, T effector cells furthermore undergo metabolic reprogramming to induce mitochondrial fusion which promotes ETC function, and this is a prerequisite for the transition from an effector to a memory cell fate (Buck et al., 2016). Furthermore, Complex III is required for the suppressive function of regulatory T cells by maintaining a regulatory gene expression profile (Weinberg et al., 2019), preventing autoimmunity by the loss of self-tolerance.

Similarly, cells of the innate immune system undergo massive metabolic reprogramming following bacterial infection. Macrophages and dendritic cells increase glycolytic flux and lactate production following microbial stimulation (O’Neill & Pearce, 2015), which is accompanied by a decrease in ETC activity in macrophages (Tannahill et al., 2012) but an increase in other cell types like dendritic cells (Everts et al., 2014). In addition to changes in ETC activity, following classical activation by LPS stimulation or interferon gamma, mouse macrophages undergo a metabolic switch that entails increased glycolysis, a broken tricarboxylic acid (TCA) cycle, and suppression of oxidative respiration (Mills et al., 2017). These metabolic changes are generally corelated with a pro-inflammatory M1 phenotype (Mills et al., 2016). One of the key drivers of this metabolic switch relies on the interferon-induced expression of inducible nitrate oxide synthase (iNOS) and the production of NO (Bailey et al., 2019), which suppresses enzymes in the TCA cycle (Palmieri et al., 2020) and ETC complexes particularly Complex I and IV (Cleeter et al., 1994; Clementi et al., 1998). The net result of this metabolic switch is the accumulation of succinate, which stabilizes HIF-1alpha to initiate a pro-inflammatory gene transcription program of classically activated macrophages (Mills et al., 2017). These observations suggests that during infections, the ETC and TCA serve functions other than sustaining ATP production for the anabolic needs of the phagocyte. Indeed, bacterial infection of murine macrophages induces architectural changes in the ETC leading to reduced CI and CI-containing supercomplexes with a concomitant rise in CII activity (Garaude et al., 2016). Such rearrangements are necessary for producing ETC ROS necessary for bactericidal activity. However, the requirement and role of ETC complexes during viral infection, and whether they undergo any form or reprogramming to modulate immune responses, remain understudied.

Another outstanding question is the role of ROS produced from specific sites of the ETC in determining cellular outcomes, particularly in the context of immune modulation. ROS production by the ETC is often considered as a homogenous entity. In reality, ETC ROS can originate from either Complex I or Complex III, and can function as messengers in cellular transduction in addition to being damaging entities in pathology. Whereas CI releases superoxide into the matrix, CIII ROS is predominantly released into the intermembrane space (IMS). Work by Dröse and colleagues suggest that while CI ROS is predominantly deleterious due to their reaction with mtDNA in the matrix, CIII-derived ROS may serve as second messengers in cellular signaling, since CIII delivers ROS into both the mitochondrial matrix and the cytosol where it can interact with cytosolic signaling pathways (Bleier & Dröse, 2013). Indeed, ROS produced from different ETC sites target distinct redox-sensitive proteins (Bleier et al., 2015). Pathogenic ROS production from CI is produced mainly by reverse electron transport at the I_Q_ site caused by elevated levels of ubiquinol or succinate and is particularly pathogenic in ischemia-reperfusion injury (Chouchani et al., 2014). CIII ROS is produced by the outer ubiquinone-binding site of complex III (CIII Qo site), which has the largest capacity of all mitochondrial sites to produce superoxide (Brand, 2016; Brand et al., 2016). CIII Qo ROS is proposed to be the main ROS species triggered by hypoxia that acts redundantly with the Von Hippau-Lindau (VHL)-prolyl hydroxylase (PH) axis to stabilize HIF-1alpha and its downstream hypoxic transcriptional program (Bell et al., 2007; Brunelle et al., 2005; Chandel et al., 2000; Klimova & Chandel, 2008). Complex III-derived ROS has also been reported to be required for adipocyte differentiation via induction of the PPARγ transcriptional machinery in an mTORC1-dependent manner (Tormos et al., 2011). In the context of immunity, CIII ROS is required for T cell activation (Sena et al., 2013), but appears to be dispensable for NLRP3 inflammasome activation (Billingham et al., 2022).

In this study, we asked if viral infection induces ETC rearrangements that function to modulate the immune response, and whether ETC CIII ROS had any roles in controlling the host response to viral infection. Severe disease in acute respiratory viral infections (ARVI), such as influenza or COVID-19, are driven by excessive production of pro-inflammatory cytokines and chemokines, which cause a hyperinflammatory phenotype that cannot be effectively treated by anti-viral therapeutics (Globenko et al., 2023; Ludwig et al., 2023). As such, there is still a large unmet need for host-targeting therapies that effectively curtail excessive host-inflammation without compromising anti-viral immunity. We therefore applied our question of ETC remodeling to Influenza A infection. Here, we show that a reduction in ETC supercomplexes and Complex III levels is an innate and host-protective response to influenza infection, leading to dampening of viral-induced inflammation and host pathology without compromising anti-viral activity. This raises the possibility that targeted suppression of Complex III might be a feasible strategy for host-targeted therapy against hyperinflammatory ARVIs.

## Results

### Viral infection of macrophages promotes respiratory chain remodeling and Complex III Suppression

To determine how ETC complex assembly is altered during viral response, we subjected murine bone marrow-derived macrophages (BMDM) to infection by mouse adapted influenza virus (A/PR8/1934 H1N1) (PR8), as well as Polyinosinic-polycytidylic acid (Poly I:C), a synthetic analog of double-stranded RNA (dsRNA) and viral mimic which induces an interferon response through Toll-like receptor 3 (TLR3). Infection of C57BL/6 mouse BMDM with PR8 results in transient non-productive infection (Figure S1A-B)(Londrigan et al., 2015) that stimulates a robust interferon response (Figure S1C). Infection and polyI:C resulted in rearrangements of the ETC as revealed by BN-PAGE analysis (Figure 1A). Specifically, respiratory supercomplexes were significantly reduced as measured by CI and CIII subunit levels in the supercomplex (SC) region. The loss of CI and CIII SCs corelated with a downward trend in OXPHOS (Figure 1B), reduced membrane potential (Figure S1D), reduced pyruvate/malate-driven CI-III-IV electron flow (Figure 1C) following PR8 and Poly I:C treatment. To understand if OXPHOS suppression is directly related to changes in the ETC, we measured enzymatic activities of Complex I and III using spectrophotometric kinetic assays. Indeed, CIII enzymatic activity was progressively reduced following both PR8 and Poly I:C exposure (Figure 1D), while CI enzymatic activity stayed unchanged or even trended upwards (Figure 1E). These experiments demonstrate that following viral infection, the ETC undergoes rapid and dynamic changes to reduce OXPHOS by reducing CIII but not CI activity. These results suggest that the primary effect of viral infection on ETC after infection might be to downregulate CIII. Since CI and CIII stability are interdependent (Acín-Pérez-Pérez et al., 2004), reduced CIII is predicted to impair CI stability, which in turn leads to the disassembly of respiratory supercomplexes observed following infection.

**Figure 1.**
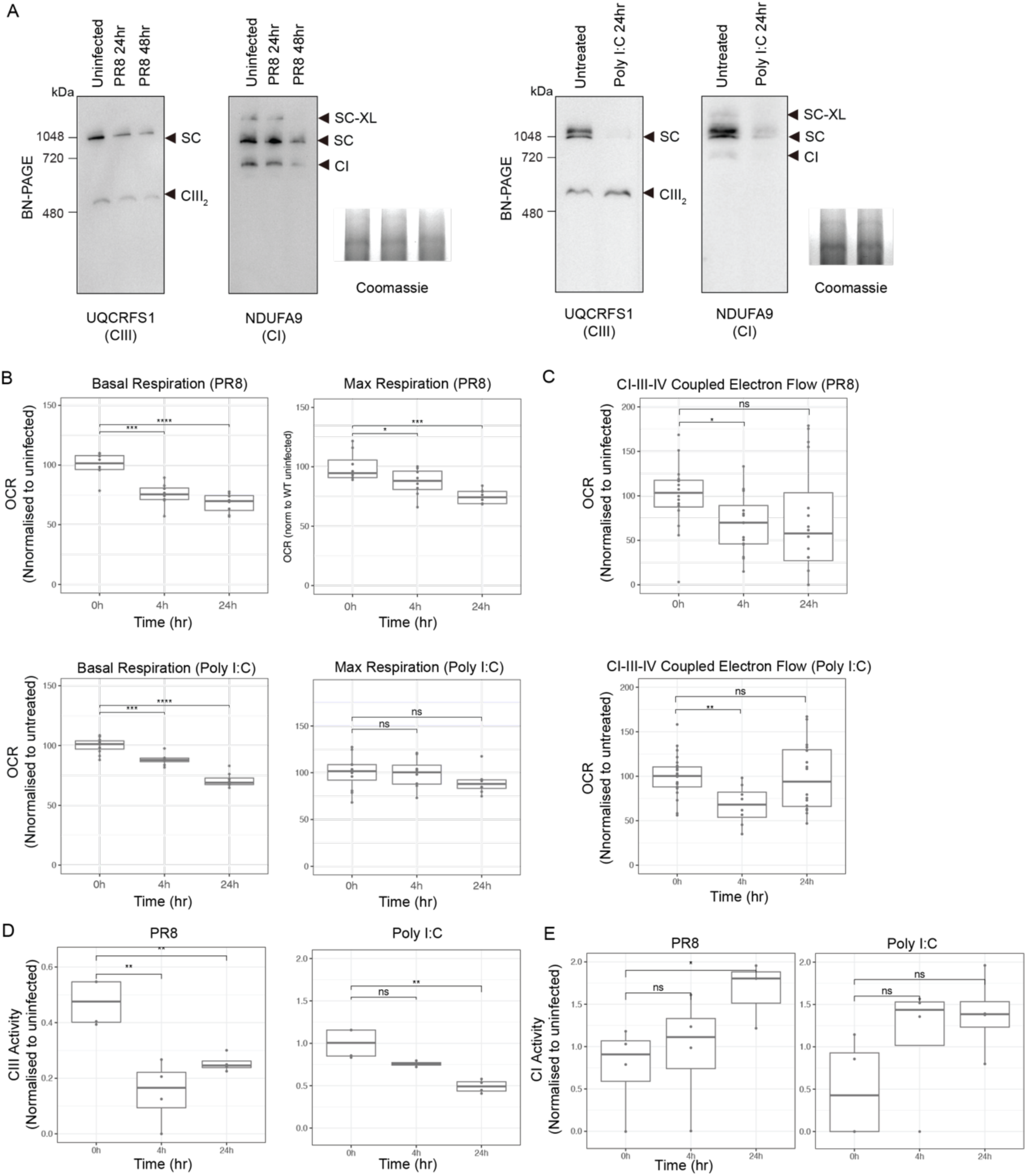
Viral infection reduces ETC Complexes and OXPHOS. (A) Blue-native (BN) PAGE of isolated mitochondria from WT BMDMs infected with PR8 (left) or treated with Poly I:C (right) harvested at indicated timepoints. Coomassie is used as loading control. (B) Seahorse oxygen consumption analysis of WT BMDMs infected with PR8 (top) or treated with Poly I:C (bottom) at indicated timepoints. Quantification of basal respiration and ATP production are shown. Values were normalised to Citrate Synthase (CS) activity and fold change was calculated over uninfected (0 hr) samples. n = 8 technical replicates of a representative experiment. (C) Pyruvate-energized CI-III-IV-coupled electron flow measured by Seahorse of WT BMDMs infected with PR8 (top) or treated with Poly I:C (bottom) at indicated timepoints. Values were normalised to CS activity and fold change was calculated over 0 hr samples. n = 8-16 technical replicates of a representative experiment. (D) Spectrophotometric respiratory chain activity assay (RCA) for CIII activity of WT BMDMs infected with PR8 (left) or treated with Poly I:C (right) at indicated timepoints. Values were normalised to CS activity and fold change was calculated over 0 hr samples. n = 4 biological replicates. P-values are from student’s t-test. (E) RCA for CI activity as per (D). BMDMs were infected with PR8 at MOI = 10 or with 10 μg/ml Poly I:C. Data are presented as mean ± SEM, n = 4 biological replicates. P-values are from student’s t-test.

### CIII activity is suppressed via downregulation of COM complexes during viral infection

To understand the mechanism responsible for CIII downregulation, we performed quantitative mass spectrometry on mitochondria isolated from PR8 and Poly I:C-treated BMDM. This analysis revealed that the most significantly downregulated mitochondrial protein 24 hours after infection or Poly I:C treatment was UQCC2 (Figure 2A). UQCC2 is an early Complex III assembly factor first characterized in yeast to be required for the translation of yeast CYTB – the enzymatic core subunit of Complex III - in the mitochondria (Gruschke et al., 2012). Likewise, in mammals, we and others recently reported that UQCC2, along with BRAWNIN (BR or UQCC6), UQCC1, SMIM4 (UQCC5) and C16ORF91 (UQCC4), form a 240 kD complex known as the <u>Co</u>-ordinator of <u>m</u>itochondrial CYT<u>B</u>, or COMB complex that is required for CIII assembly via the stabilization of nascent CYTB (Dennerlein et al., 2021; Liang et al., 2022). We confirmed the downregulation of UQCC2 and its client protein CYTB by western blotting (Figure 2B). BR, a core stabilizing subunit of the COMB complex was also significantly reduced by SDS-PAGE (Figure 2B) as well as on native PAGE (Figure 2C), where BR runs at 240 kD marking the position of the COMB complex. Consequently, the COMB complex – as marked by UQCC1 (Figure 2C) (Liang et al., 2022) - was significantly reduced following infection and Poly I:C. The related COMA complex, which contains UQCC1 and 2 but not BR and mediates the exit of CYTB from the mitochondrial ribosome exit tunnel was also reduced (Figure 2C). This suggests that during a viral-induced interferon response, macrophages downregulate BR and UQCC2 leading to reduced COMA/B complexes and CYTB production. Consequently, CIII assembly and OXPHOS are depressed following infection.

**Figure 2.**
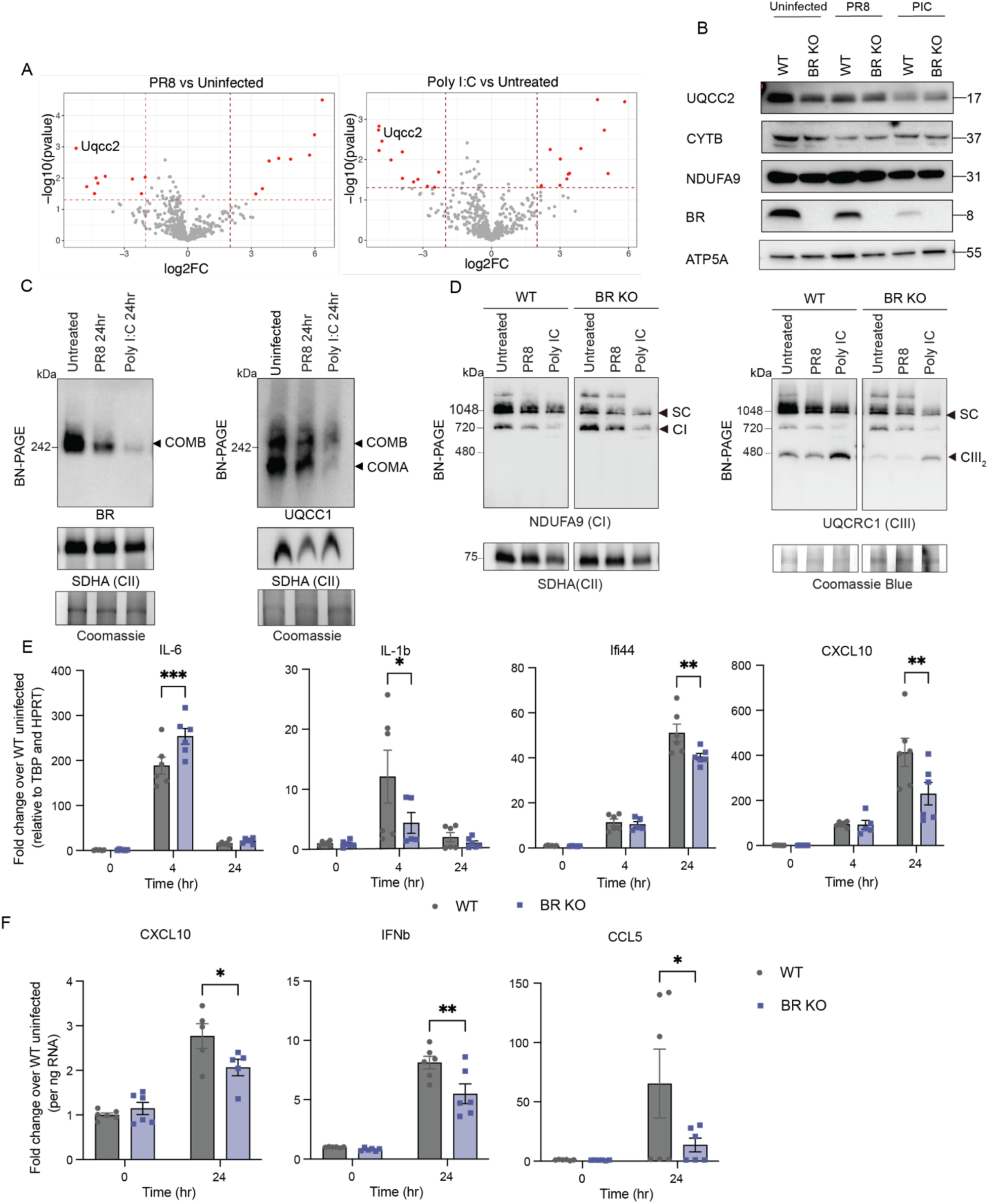
Complex III downregulation is an innate response following viral infection and dampens the immune response. (A) Label-free quantitative proteome analysis of mitochondria isolated from WT BMDMs infected with PR8 (left) or treated with Poly I:C (right) for 24 hr. Mean values of log_2_ fold change (treated / untreated) for each detected protein are derived from 3 biological replicates. (B) SDS PAGE of mitochondria isolated from WT BMDMs infected with PR8 or Poly I:C for 24 hr. (C) BN PAGE of mitochondria isolated from WT BMDMs infected with PR8 or treated with Poly I:C at indicated timepoints. (D) BN PAGE of mitochondria isolated from WT and BR KO BMDMs infected with PR8 or treated with Poly I:C for 24 hr. (E) Interferon stimulated gene expression in WT and BR KO BMDMs following PR8 infection at indicated timepoints. Fold change was calculated over WT 0 hr samples. (F) Pro-inflammatory cytokines secretion by WT and BR KO BMDMs following PR8 infection at indicated timepoints. Fold change was calculated over WT 0 hr samples. BMDMs were infected with PR8 at MOI = 10 or with 10 μg/ml Poly I:C. Data are presented as mean ± SEM, n = 5 biological replicates. P-values are from student’s t-test.

### Reduction in CIII assembly via BR knockout results in an anti-inflammatory phenotype

We next asked if the observed reduction in CIII following infection is an adaptive or maladaptive phenotype. To do this, we examined BMDMs from BR KO mice, which are deficient in COMB complexes, and have reduced levels of CIII at steady state (Liang et al., 2022) (Figure 2D). Following Pr8 infection, BR KO BMDMs displayed a lower level of pro-inflammatory and interferon-induced gene expression as exemplified by *IL1β, Ifi44, Cxcl10* following PR8 infection (Figure 2E). BR KO BMDMs also secreted significantly less proinflammatory cytokines CCL5, CXCL10 and IFNΒβ following PR8 infection (Figure 2F). Interestingly, IL-6 expression was upregulated by PR8 infection in BR KO BMDMs (Figure 2E), despite the general downregulation of other pro-inflammatory cytokines. IL-6 has previously been shown to protect against H1N1 infection by sustaining neutrophil-mediated clearance of infected cells in the lungs (Dienz et al., 2012), even though it has also been implicated in the pathogenesis of other SARS-CoV2-mediatd respiratory infection (Blanco-Melo et al., 2020). This raised the question of whether BR KO might be better protected against H1N1 infections *in vivo*. Since PR8 infection in C57BL6-derived BMDMs is non-productive, the observed anti-inflammatory effects of BR KO could either be due to reduced infectivity of Br KO cells or to a reduced inflammatory response triggered by the infection. We ruled out reduced infectivity by quantifying intracellular viral particles right after the infection (0hr), which showed no differences between WT and BR KO BMDMs (Figure S2A). To confirm this, we repeated the experiment in WT and BR KO MLE-12 cells, which are murine lung epithelial cells that support productive infection by PR8. Similar to BMDMs, BR KO MLE-12 had lower levels of pro-inflammatory cytokine gene expression (Figure S2B) and secretion (Figure S2C) even though viral copy number was unchanged or even higher in Br KO cells (Figure S2D). Altogether, our data suggest that BR KO BMDMs mount a less proinflammatory response following viral infection that is independent of viral replication.

### Inhibition of Complex III ROS dampens inflammation in viral-infected BMDMs

Despite having reduced levels of CIII activity (Figure S3A), Br KO BMDMs had less OXPHOS suppression following PR8 infection (Figure 3A), in line with the observed reduction in pro-inflammatory response. These data suggest a change in metabolic reprogramming downstream of CIII suppression in Br KO that confers an anti-inflammatory effect. We next sought to define the mechanism downstream of CIII suppression that results in reduced inflammation. Reduced CIII activity has 3 major effects: i. increased reduced Coenzyme Q (ubiquinone) and QH_2_/Q ratio; ii. accumulation of succinate; and iii. reduced ROS production from the CIII Qo site, which is the main source of mitochondrial ROS production. While increased QH_2_/Q and succinate have been associated with enhanced macrophage inflammation (Mills et al., 2017) and thus unlikely to explain suppressed inflammatory phenotype of BR KO, the role of CIII Qo site in inflammation has not been explored. Consistent with reduced CIII activity, we found that CIII Qo site ROS was markedly reduced in BR KO BMDMs at basal, as well as 24h post-infection (Figure 3B), but not ROS produced from CI either at the flavin or Q binding sites (Figure S3B). Reduction in CIII Q_o_ ROS in BR KO BMDM resulted in an overall decrease in mitochondrial ROS as measured by MitoSOX without a change in mitochondrial membrane potential (Figure S3C). These results suggest that CIII Q_o_ site ROS is the major contributor of ROS during viral infection and might potentiate the innate inflammatory response. To test this, we treated BMDMs with S3QEL, a small molecule that binds to the CIII Q_o_ site to reduce CIII ROS without inhibiting the forward flow of electrons and OXPHOS activity (Orr et al., 2015). Consistent with the hypothesis that CIII Q_o_ ROS promotes inflammation during viral infection, BMDMs pre-treated with S3QEL had lower levels of inflammatory gene expression following PR8 infection (Figure 3C). S3QEL treatment also phenocopied the trend towards increased *IL6* expression as seen in Br KO BMDMs (Figure 3C). We next sought to ask what the downstream mechanism of CIII ROS in controlling inflammatory phenotype might be. CIII ROS has been shown to induce HIF1/2 stabilization cardiac muscle (Guzy et al., 2005). In macrophages, HIF1/2 stabilization following TLR stimulation activates aerobic glycolysis, inflammatory cytokines expression, as well as inducible nitric oxide synthase (NOS2) to reinforce macrophage activation (Knight & Stanley, 2019). However, while S3QEL suppressed *Nos2* in BMDMs following LPS stimulation (Figure S3D), it had no effect following PR8 infection (Figure S3E). Hence, this is unlikely the mechanism involved in the observed dampened inflammation. A second possible effect of CIII ROS is to inhibit fatty acid oxidation (FAO). CIII ROS has previously been shown to target proteins in the fatty acid oxidation (FAO) pathway to inactivate the enzymes (Bleier et al., 2015). Increased FAO is corelated to alternative activation (M2) of macrophages, which are less pro-inflammatory (Liu et al., 2017). Indeed, following PR8 infection, Br KO BMDMs have a higher rate of FAO compared to WT BMDM (Figure 3D). Lastly, treatment of PR8-infected BMDMs with the FAO inhibitor etomoxir (ETO) was sufficient to increase pro-inflammatory cytokine expression (Figure 3E). Therefore, CIII ROS likely downregulates FAO to increase proinflammatory macrophage polarization following viral infection. These results suggest that CIII suppression following infection might serve to curtail the production of CIII ROS, and to reduce the proinflammatory response as an intrinsic mechanism to resolve or dampen inflammation.

**Figure 3.**
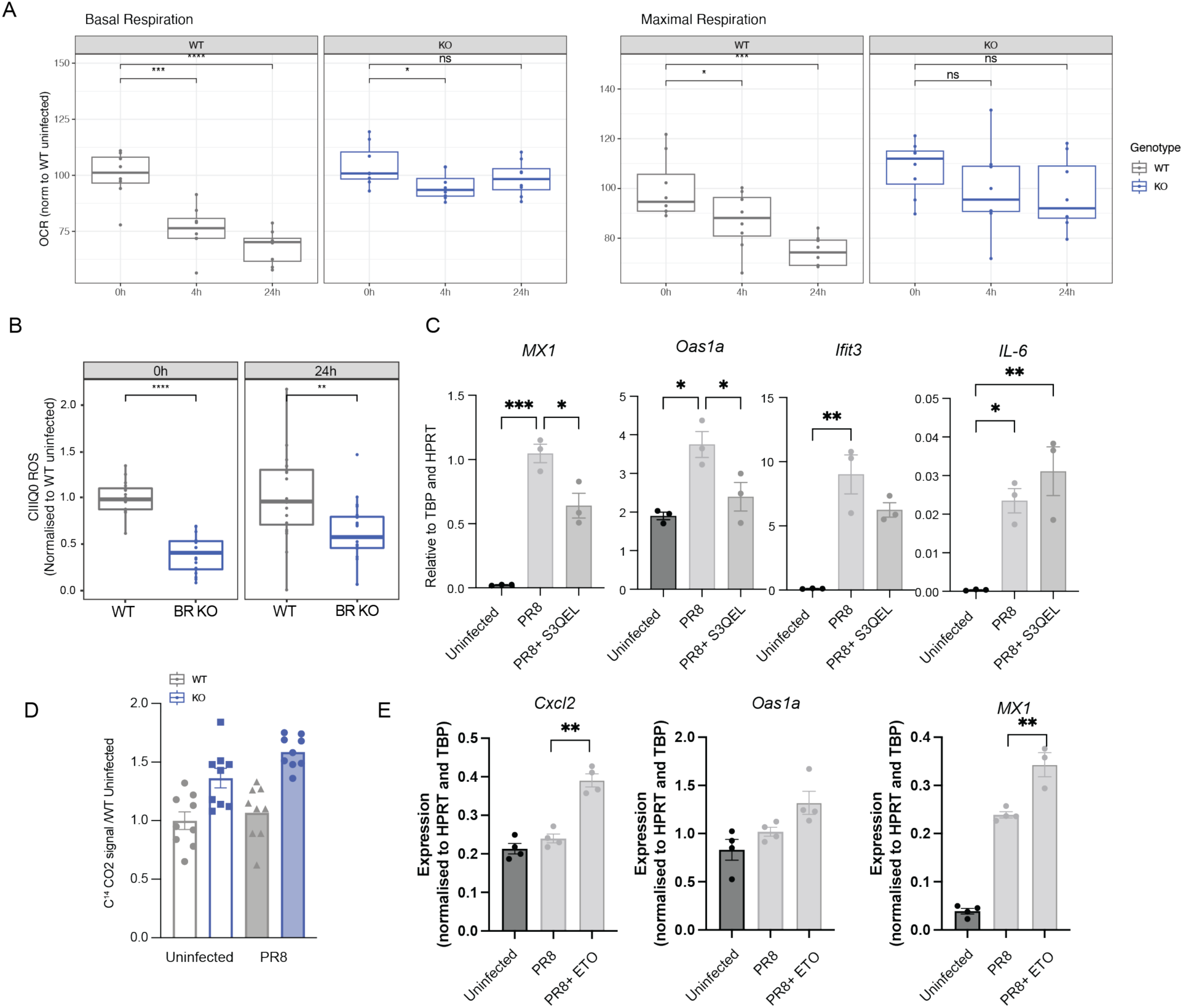
Infection dampens CIII activity and results in reduced mitochondrial ROS generation associated with a less pro-inflammatory profile. (A) Seahorse oxygen consumption analysis of WT and KO BMDMs infected with PR8 at indicated timepoints. Quantification of basal and maximal respiration are shown. Values were normalised to Citrate Synthase (CS) activity and fold change was calculated over uninfected (0 hr) samples. n = 8 technical replicates of one representative experiment. (B) CIII Q_o_ site-mediated ROS generation in WT and BR KO BMDMs infected with PR8. Values were normalised CS activity and then to WT 0 hr samples. n = 8 technical replicates of a representative experiment. (C) Interferon stimulated gene expression in WT and BR KO BMDMs treated with 5 μM S3QEL following PR8 infection (MOI = 10). n = 3 biological replicates. (D) Radioactive palmitate oxidation assay to measure FAO rate in WT and BR KO BMDMs following PR8 infection (MOI=10). n = 9 biological replicates (E) Interferon stimulated gene expression in WT and BR KO BMDMs treated with 5 μM Etomoxir following PR8 infection (MOI = 10). n = 4 biological replicates.

### CIII reduction is protective against mouse IAV infection

We next sought to determine how CIII reduction *in vivo* affects the outcome of an infection *in vivo*. Severe influenza infection is associated with excessive host inflammation and cytokine storm. We hypothesized that CIII reduction in Br KO mice would dampen host inflammation during influenza infection. At the basal uninfected state, WT and Br KO lungs have no visible differences histologically (Figure S4A) or in the composition of leucocytes in bronchial alveolar fluid (BALF) (Figure S4B). Following a physiological dose of PR8 (250 PFU), Br KO mice had reduced weight loss and recovered more quickly compared to WT mice (Figure 4A). Pathological examination showed that Br KO mice had almost no externally visible lung lesions compared to WT mice (Figure 4B) at 5 days post-infection (dpi). Histology of lung sections revealed reduced leucocytic infiltration into the lung parenchyma, alveolar wall and spaces, with reduced thickening of the bronchial epithelium and bronchiolar expansion in Br KO mice following PR8 infection (Figure 4C). These observations suggest that CIII reduction attenuates immune activation during PR8 infection. Consistent with the histologic phenotype, lung homogenates of Br KO mice contained lower levels of pro-inflammatory cytokines such as IL-6, Tnf-ɑ, IL-1ɑ and Ccl2 (Mcp-1) (Figure 4D). We next asked if Br KO would be sufficient to protect mice against a lethal influenza infection. Following a challenge of 1250 PFU with PR8, which causes severe lethal infection in mice, Br KO mice did not have improved overall survival (Figure S4C). However, 2/12 (17%) of Br KO mice that survived the high-dose challenge regained body mass more rapidly than surviving WT mice (2/13 or 15%) (Figure S4D), consistent with the finding that KO mice sustain less organ damage induced by PR8 infection. Hence, in the context of severe lethal infection, CIII suppression alone is insufficient to reduce mortality.

**Figure 4.**
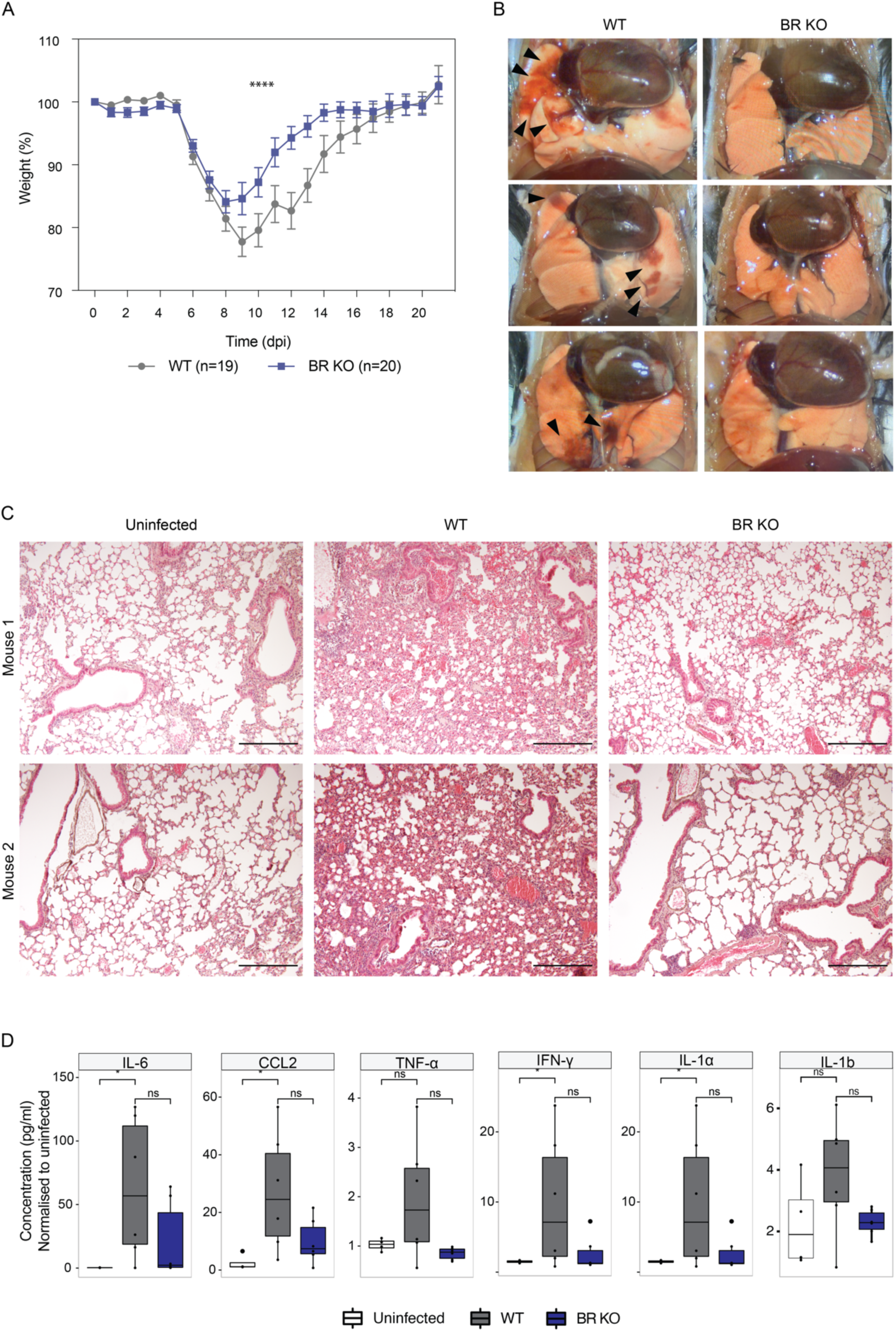
CIII suppression protects mice from H1N1 (PR8) infection. (A) Body weights of WT and BR KO mice inoculated with 250 PFU of PR8 as a percentage of initial body weight before infection. Data is presented as mean ± SEM, n = 19 (WT) and 20 (BR KO). P-value is from Anova analysis. (B) Representative images of gross pathology of WT and BR KO mouse lung inoculated with 250 PFU of PR8 at 5 dpi. Black arrowheads indicate lesions due to infection. (C) Representative images of alveolar pathology and leucocyte infiltration visualised by H&E staining of WT and BR KO mouse lung inoculated with 250 PFU PR8 at 3dpi. Scalebar = 5um. (D) Pro-inflammatory cytokine (IL-6, CCL2, TNFα, IFNγ, IL-1α, and IL-1β shown) levels in lung homogenates of WT and BR KO mice following PR8 infection at 3dpi. Fold change was calculated over WT uninfected mice. Data is presented as mean ± SEM, n = 6 mice.

### CIII reduction *in vivo* is associated with reduced pro-inflammatory monocytes

To understand the possible mechanisms of reduced host inflammation in Br KO mice, we performed single-cell RNAseq of mice infected with a low (250 PFU) and high (1250 PFU) dose of PR8. This analysis identified 19 distinct cell types of epithelial, endothelial and leucocytic subtypes (Figure S5A) and showed that there were no differences between WT and KO at the basal uninfected state (Figure S5B). Of note, the monocytic populations could be further clustered into 3 subtypes (Figure S5C): classical monocytes, proinflammatory classical monocytes (called inflammatory monocytes hereafter) and non-classical monocytes or macrophages (Mono /Macs) (Figure S5C). Inflammatory monocytes were distinguished by high expression of *Ly6c*, *FcgRI*, *Isg15* expression (Figure S5C,E), as well as signatures of proinflammatory pathways such as interferon beta-1 and TNF-alpha (Figure S5F). ViralTrack (Bost 2020) analysis revealed that inflammatory monocytes were the predominant cell types in which PR8 viral genomes could be detected (Figure 5A). At 250 PFU, PR8 viral genomes were not detected in Br KO cells by scRNAseq at all (Figure 5A), whereas at 1250 PFU, Br KO leucocytes had reduced viral sequences detected compared to WT cells (Figure 5A). Consistent with these ViralTrack results, qPCR quantification of PR8 viral genome in lung homogenates at 5dpi confirmed that Br KO lungs had lower viral copy number compared to WT (Figure 5B). These data indicate that despite reduced host inflammation in Br KO mice, antiviral resistance was not compromised, and in fact Br KO animals were better able to control viremia compared to WT mice. To ascertain if there were changes in the frequency of leucocytes recruited following PR8 infection in WT and KO mice, we performed Milo analysis, a statistical framework that performs differential abundance testing of cell populations between conditions (Figure 5C) (Dann 2022). This analysis revealed that at 3dpi, 250PFU-infected Br KO lungs had significantly reduced numbers of inflammatory monocytes. 1250 PFU-infected Br KO lungs showed a similar trend, in addition to reduced numbers of dendritic cells and classical monocytes (Figure 5D-E). Using high expression of Ly6c as gating strategy to define inflammatory monocytes (Figure S5D), flow cytometric analysis confirmed the observed reduction in the number of Ly6c^hi^ inflammatory monocytes (CD45^+^,Ly6G^-^,Ly6c^hi^,Cd11b^+^,Cd11c^-^) in Br KO lungs at 5dpi (Figure 5F, see Methods for gating strategy). This population of inflammatory monocytes are transiently induced or recruited following PR8 infection, since they are absent at 0-1dpi, increased at 3dpi and reduced by 5dpi (Figure S5D). Inflammatory monocytes have been shown to contribute to severe influenza in both mice and humans (Lin 2014,Coates 2018,Cole 2017). Consistent with this hypothesis, inflammatory monocytes are enriched in signatures of interferon activity and inflammatory response, IL6-JAK/STAT signaling as well as the monocytic signature of hyperinflammatory monocytes found in severe SARs-COV2 infection (Bost 2020) and severe influenza in juvenile mice (Figure S5F-G). The peak of the inflammatory response in these monocytes appear to be at 3dpi (Figure S5G), consistent with previous studies demonstrating that proinflammatory factors in H1N1 are produced in waves by different populations of myeloid cells (Zhang 2020). Importantly, the inflammatory signatures associated with severe flu and COVID, as well as inflammasome activity and TNFa/interferon signaling were significantly reduced in Br KO inflammatory monocytes (Figure 5G), consistent with their reduced numbers and overall reduction in inflammatory cytokine production in Br KO lungs. To assess if the metabolic changes detected in BMDMs were also seen in monocytes *in vivo*, we utilized Compass (Wagner et al., 2021), an algorithm to infer cellular metabolic states utilizing single-cell RNA sequencing and flux balance analysis. Consistent with reduced inflammation and the in vitro studies in BMDMs, Br KO inflammatory monocytes were predicted by Compass to have increased flux through the TCA cycle (Figure S6A) and FAO pathway (Figure S6B). In line with these observations, Br KO inflammatory monocytes also showed a significantly reduced M1-like inflammatory score compared to WT (Figure S6C), alongside an increase in FAO score (Figure S6D). In line with this, Monocle pseudotime analysis (Figure S6E-F) positioned WT inflammatory monocytes later along an inflammation-associated trajectory (Figure S6G), whereas Br KO cells remained earlier in pseudotime (Figure S6I), indicating attenuated progression toward a more highly inflammatory state. These results suggest that CIII suppression in these monocytes might lead to reduced metabolic switching that underlies the dampened inflammatory response. The reduction in inflammatory status was however not limited to this monocytic population. Instead, GSEA analysis indicates that virtually all myeloid populations had reduced cytokine and interferon response, reduced viral defense signatures and increased transcription related to cytosolic translation, possibly reflecting secondary effects of reduced viral infection or reduction in pro-inflammatory monocyte activity (Figure 5J). The overall outcome is host protection against IAV-induced lung inflammation that does not compromise host anti-viral defense as evidenced by lower viral copy number in Br KO animals.

**Figure 5.**
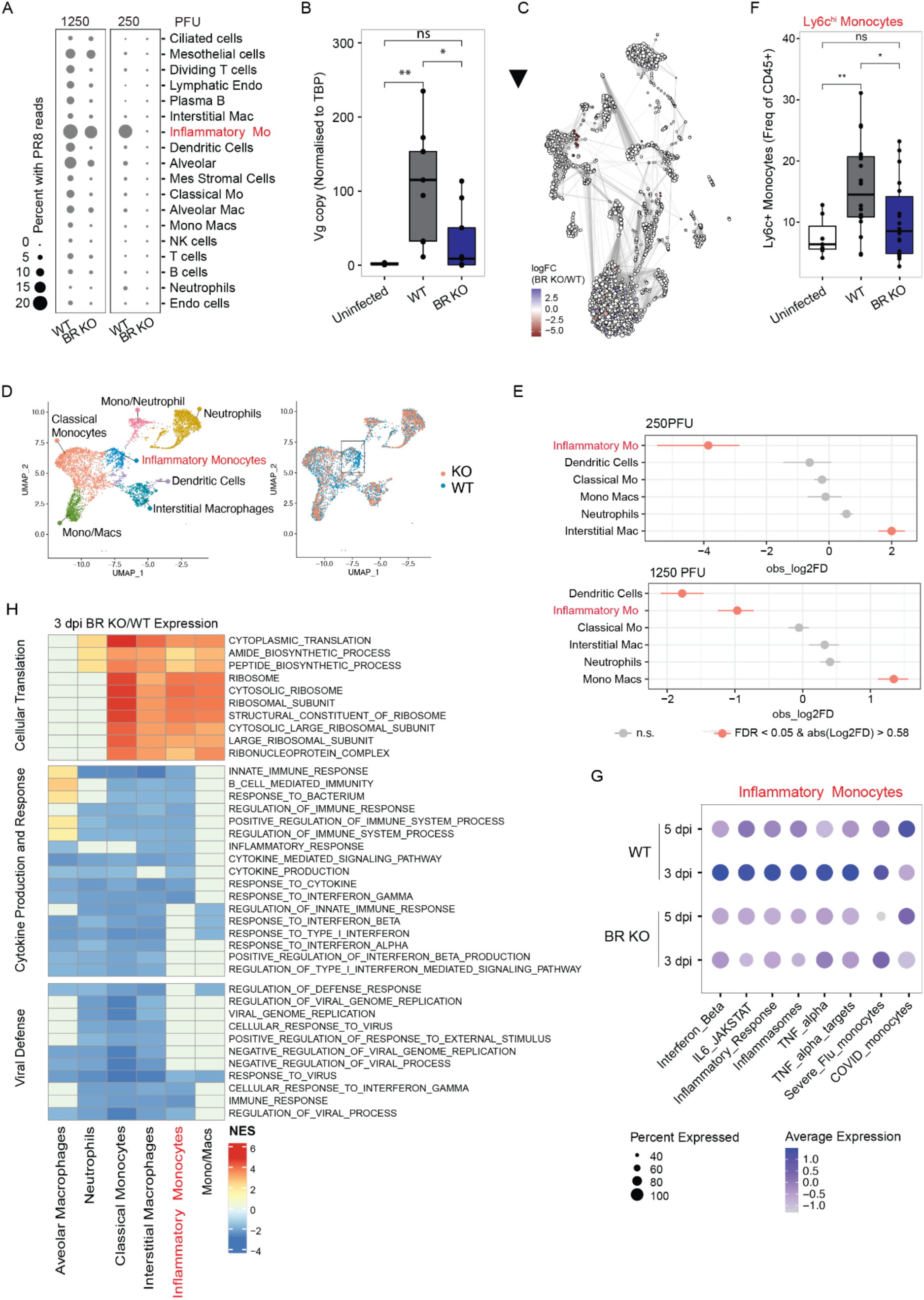
CIII suppression reduces inflammatory monocyte and inflammatory lung phenotype following H1N1 (PR8) infection. (A) Viral track analysis of WT and BR KO lungs infected with PR8 at indicated doses at 3dpi, n = 3. (B) Viral genome copy number in WT and BR KO lungs infected with PR8 (250 PFU) at 3 dpi quantified by qPCR. Data is presented as mean ± SEM, n = 6. (C) MiloR analysis for differential cell abundances in WT and BR KO lungs infected with PR8 (250 PFU). (D) Umap of myeloid cell populations highlighting the abundance of inflammatory monocytes (boxed) in WT versus KO lungs at 3 dpi after PR8 (250 PFU) infection. (E) Quantitative analysis of differential abundance of myeloid cell populations in (D). (E) Flow cytometry staining of mouse Ly6C+ monocytes isolated from WT and BR KO lungs infected with PR8 (250 PFU) at 3dpi. Data is presented as mean ± SEM, n = 15. (F) GSEA analysis of WT and BR KO lungs infected with PR8 (250 PFU) at 3dpi reveal reduced expression of genes involved in inflammation and viral defence in BR KO mice. (G) Differential expression of the indicated gene signatures derived from MSigDB in WT vs KO inflammatory monocytes at after PR8 (250 PFU) infection.

### CIII suppression reduces monocyte migration

To further define a possible reason for the reduced numbers of inflammatory monocytes in BR KO, we examined intercellular interactions in the PR8-infected lung using CellChat (Jin et al., 2021), focusing on incoming and outgoing signals of this population (Figure 6A). While there were no dramatic alternations in overall predicted intercellular communications, CellChat predicted a reduced level of CCL chemokine-mediated signaling in BR KO inflammatory monocytes (Figure 6A). Specifically, CellChat predicted that CCL2-CCR2 signaling between inflammatory monocytes and various cell types such as macrophages, other monocyte subsets and neutrophils was absent in BR KO (Figure 6B). Closer examination revealed that this prediction was driven primarily by a lack of induction of the chemokine Ccl2 (Mcp-1) in Br KO leucocytes (particularly monocytes) (Figure 6C) following infection. Ccl2 is is a key mediator of inflammatory monocytic migration signaling through the C-C chemokine receptor Ccr2 (Boring et al., 1997). In order to validate the Cell Chat prediction that Br KO monocytes might have reduced migratory capacity, we performed an *in vitro* migration assay wherein primary bone-marrow derived monocytes were seeded into a transwell insert and allowed to migrate for 5 hours towards PR8 infected MLE-12 lung epithelial cells or BMDM macrophages (Figure 6D). This analysis revealed that BR KO monocytes indeed had reduced migration compared to WT monocytes (Figure 6D). Of note, WT monocytes migrated equivalently towards PR8-infected WT or KO MLE-12 or BMDMs (Figure S7A), suggesting that the defects in migration are cell-intrinsic to BR KO monocytes. Lastly, in line with the prediction that CIII ROS contributes to the pro-inflammatory phenotype of infected monocytes, pre-treatment of PR8-infected BMDMs with S3QEL reduced migration (detached cells) (Figure 6F). Altogether, CIII reduction and the ensuing reduction of CIII ROS promotes both host tolerance and host anti-viral defense during H1N1 infection, partially by modulating the abundance or activity of Ly6c^high^ monocytes that are induced following influenza infection. This is in line with a recent study demonstrating that an IFN-gamma regulated subset of Ccr2 + monocytes drives lung damage during influenza infection (Schmit et al., 2022). Ablation of this population through the genetic deletion of Ccr2 or IFN-gamma reduced lung inflammation, pathology, and disease severity without impacting viral clearance (Schmit et al., 2022).

**Figure 6.**
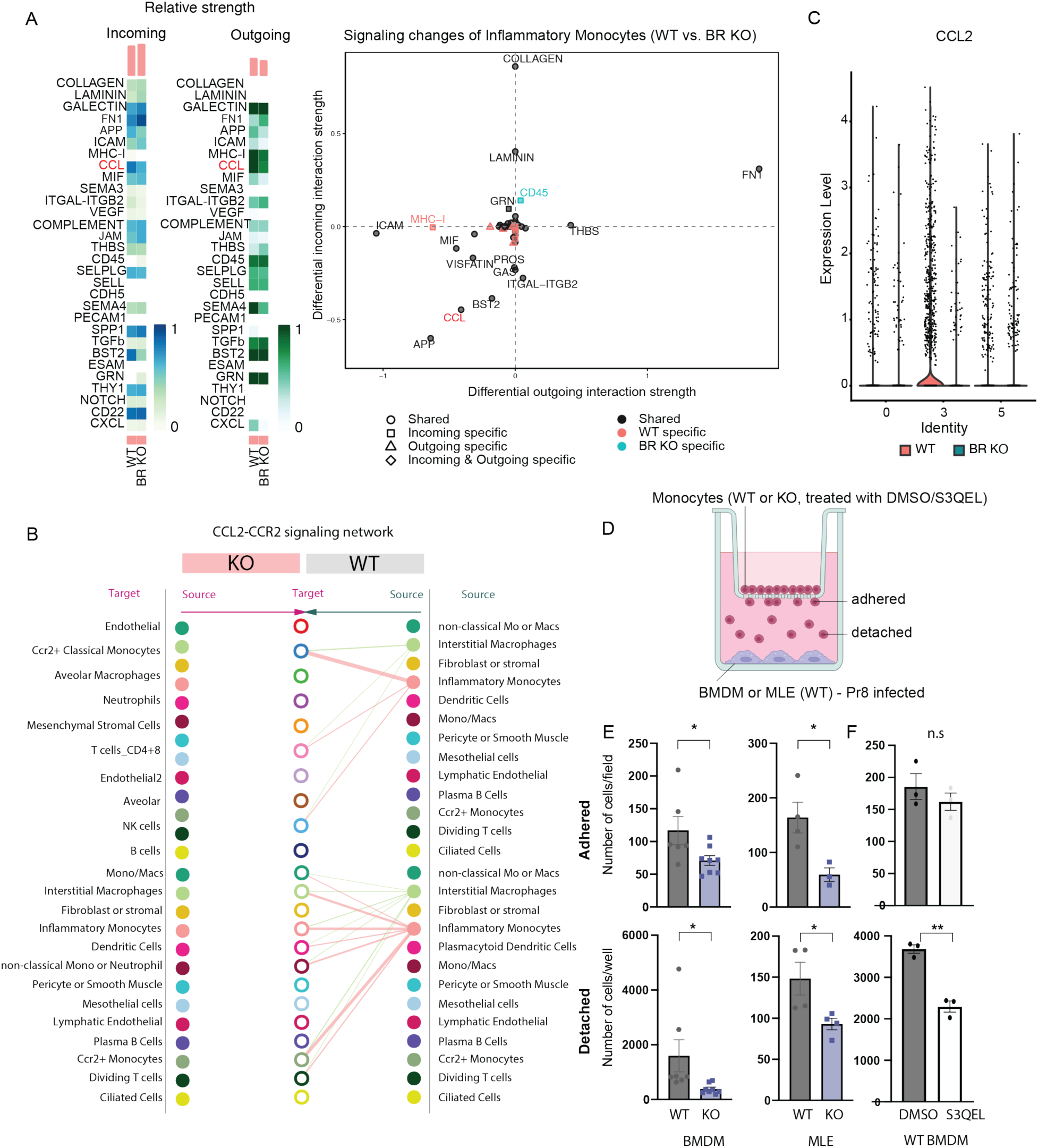
BR KO monocytes have reduced propensity to migrate in response to infection cues and exhibit reduced CCL2-CCR2 signalling. (A) Cell chat analysis of WT and BR KO lungs infected with PR8 (250 PFU) at 3dpi and 5 dpi, n = 3. Heatmaps indicate the relative strengths of interactions (left) and the plot on the left (B) Cell chat analysis reveal significantly dampened Ccl2-Ccr2 signalling axis in BR KO lungs infected with PR8 (250PFU). Data shown is a combination of expression at 3dpi and 5 dpi, n = 3. (C) *Ccl2* (*Mcp-1*) expression in WT and BR KO lungs infected with PR8 (250 PFU) at indicated timepoints (dpi), n = 3. (D) Schematic of monocyte migration assay used in (E). (E) WT and BR KO monocyte migration in response to BMDMs (left) or MLE-12 (right) infected with PR8 (MOI = 10 or 1, respectively). BMDMs were infected for 8 hr before the monocyte transwells were inserted and allowed to migrate for an additional 5 hr. Migrated adherent monocytes (top) were visualised and quantified with Hoescht staining whereas migrated detached monocytes (bottom) were quantified by flow cytometry. Data are presented as mean ± SEM, n = 4-6.

## Discussion

In this study, we provide evidence that a reduction in CIII levels via repression of the COMB assembly complexes is an innate immune response to viral infection. Genetic dampening of CIII activity in COMB-deficient BR KO mice results in reduction of mitochondrial ROS originating from CIII, and is corelated with reduced metabolic switch (i.e. OXPHOS suppression) following infection. Consequently, inflammatory responses are dampened. In vivo, CIII suppression confers protection against influenza H1N1-induced inflammation by reducing the numbers of inflammatory monocytes that contribute to hyperinflammatory disease. This reduction might be caused by reduced migration of monocytes in response to viral PAMPs or chemokines originating from infected cells. Our results show that CIII activity is required for maximal immune response following infection, suggesting that CIII reduction after activation could be part of the pro-resolving phase.

There is a paradox arising from the observation that the BR KO mice have lower viral load despite mounting a weaker immune response. Our in vitro BMDM and MLE12 models show reduced inflammation without a reduction in viral titer, suggesting an uncoupling of the anti-inflammatory effects of CIII suppression from viral load. However, our *in vivo* infection model showed both reduced inflammation and reduced viral load. As such, we cannot exclude the possibility that reduced inflammation *in vivo* is secondary to reduced viral replication or infection. Nonetheless, we speculate that reduced CIII ROS from the Qo site leads to reduced ROS-mediated signaling in monocytes, impairing their activation and recruitment into the lung parenchyma. To understand the mechanism of CIII suppression in immune dampening, it would be necessary to define the targets of CIII ROS that influence monocyte migration, activation and inflammatory profile. Lastly, we cannot exclude the possibility that non-immune cells, such as lung epithelial cells (Figure S2C-D) could also be contributing to the observed effects, since our study used a whole body BR KO. A myeloid specific KO will be necessary to clarify this point in the future.

In summary, our study demonstrates that suppression of ETC Complex III is an adaptive innate immune response that helps control excessive inflammation during viral infections, without compromising antiviral defense. This suggests that targeted modulation of Complex III activity could be a potential host-directed therapy for acute respiratory viral infections with a hyperinflammatory component. This study should motivate investigations aimed at assessing the potential of pharmacological agents such as S3QEL as viable host-directed therapeutic options.

## Acknowledgements

We thank the Duke-NUS flow cytometry core for its assistance. This work was supported by MOE-T2EP30122-0011 awarded to LH, and MOE-000095-01 awarded to ASJ. CL was supported by MOH-OFYIRG20nov-0009.

## Methods

### Cell Culture

MLE 12 cells (American Type Culture Collection, CRL-2110) were cultured using standard tissue culture techniques in Dulbecco’s Modified Eagle’s Mediusm (DMEM) : Ham’s F/12, 1:1 mix media (Lonza, 12719F) supplemented with 10% fetal bovine serum (FBS) (HyClone, SV30160.03HI) and 1% penicillin-streptomycin (Hyclone, SV30010). Cells were maintained in a humidified 5% carbon dioxide atmosphere at 37 °C.

### Isolation of Mouse BMDMs

Mouse bone marrow-derived macrophages (BMDMs) were isolated and derived using reference protocols. Briefly, bone marrow cells were harvested by flushing sterilised femurs and tibias of BR WT and KO C57BL/6 mice, followed by erythrocyte lysis using a commercial lysis buffer (BioLegend, 420301). Primary macrophages were subsequently derived from the bone marrow cells by culturing for 7 days in macrophage differentiation media (DMEM (Hyclone, SH30022.010) supplemented with 20% L929 cell-conditioned media, 10% FBS, penicillin-streptomycin, MEM non-essential amino acid solution (Gibco, 11140050), 1 mM sodium pyruvate (Gibco, 11360070), and 25 μM 2-mercaptoenthanol (Gibco, 21985023)). Cells were maintained in a humidified 5% carbon dioxide atmosphere at 37 °C with culture medium replaced on day 3 to select for adherent cells.

L929 cell-conditioned media was prepared by culturing L929 cells at 37 °C in DMEM supplemented with 10% FBS, 2 mM glutamine (Gibco, 25030081), and penicillin-streptomycin to confluence before replacing culture media and maintaining the cells at 32 °C for 10 days. L929 cell-conditioned media containing macrophage colony-stimulating factor was subsequently harvested and passed through a 0.2 μm filter (PALL Life Science, PN4612).

### In vitro Influenza Infection

Mouse-adapted Influenza H1N1 virus A/PR/8/34 (PR8) was obtained from AVS Bio and the use of the virus in vitro was approved by and carried out in accordance with BSL2 safety regulations. Cultured BMDMs or MLE 12 cells were inoculated with the PR8 virus at a multiplicity of infection (MOI) of 10 for 2 hours before culture media was washed off and replaced. Cells were returned to the incubator for infection to persist until the time of harvest or assay. Untreated control cells were given media.

### In vitro Poly I:C Treatment

Polyinosinic:polycytidylic acid (Poly I:C) (Sigma-Aldrich, P9582), is a viral mimic used to invoke immune activation of BMDMs. Cells were treated with Poly I:C at a final well concentration of 10 or 20 μg/ml and returned to the incubator until the time of harvest or assay. Untreated control cells were given media.

### RNA Extraction

RNA from BMDMs was isolated using the Direct-zol RNA Miniprep kit (Zymol, R2050) according to the manufacturer’s instructions. Cells were seeded in 24-well plates at a density of 250,000 cells per well 1 day prior to the experiment. Cell lysate was subsequently harvested at the respective timepoints after infection or treatment using the cell lysis buffer provided by the kit.

Total RNA from mouse lung tissue was isolated by first performing a homogenisation step. Minced lung tissue suspended in 1ml of TRIzol (Invitrogen, 15-596-018) was homogenised using 1 mm zirconia beads (BioSpec, 11079110) in an automatic sample processor (Roche, MagNA Lyser). Samples were pelleted to remove any remaining tissue and an equal volume of ethyl ethanol (Sigma-Aldrich, E7023) was added before being loaded on to RNA extraction columns from the Direct-zol RNA Miniprep kit.

### Interferon-stimulated Gene Expression

Isolated mRNA was reversed transcribed into cDNA using iScript Reverse Transcription Supermix (Bio-Rad Laboratories, 1708841) in a C1000 Touch Thermal Cycler (Bio-Rad Laboratories, 1851148). Quantitative PCR (qPCR) was performed using iTaq Universal SYBR Green Supermix (Bio-Rad Laboratories, 1725124) with gene-specific primers (Integrated DNA Technologies) in a CFX384 Touch Real-Time PCR Detection System (Bio-Rad Laboratories, 1855484). Primer sequences used are listed in Supplementary Table 1.

### Seahorse MitoStress Assay/Metabolic Assays

BMDM oxygen consumption rate was measured using the Agilent’s Seahorse platform according to the manufacturer’s instructions. Briefly, BMDMs were seeded on Seahorse XF96 Cell Culture Microplates (Agilent, 101085-004) at a density of 60,000 cells per well 1 day prior to the assay. Additionally, the XFe96 sensor cartridge was hydrated with distilled water and incubated at 37 °C in the absence of CO_2_ overnight. On the day of the assay, cells were washed once and media was replaced with XF basal DMEM (Agilent, 103575-100) supplemented with 1 mM pyruvate (Agilent, 103578-100), 2 mM glutamine (Agilent, 103579-100), and 10 mM glucose (Agilent, 103577-100). The pre-hydrated sensor cartridge was calibrated with Seahorse XF Calibrant (Agilent, 100840-000) for 1 hr prior to the loading of drug ports of the sensor cartridge. Both the cell culture microplate and the sensor cartridge were incubated at 37 °C in the absence of CO_2_ for at least 45 mins before the start of the assay. Drug injections for the Agilent Seahorse MitoStress assay are as follows, 2 μM of oligomycin (Sigma-Aldrich, 75351), 1 i of FCCP (Sigma-Aldrich, C2920), and 1 μM of Rotenone (Sigma-Aldrich, R8875)/Antimycin (Sigma-Aldrich, A8674). Data was obtained using the XF96 Seahorse wave software (Agilent) and normalised to citrate synthase activity.

### Citrate Synthase Activity Assay

Citrate synthase activity was quantified and used for normalisation of Seahorse MitoStress data. At the end of the Seahorse MitoStress assay, remaining media was aspirated, and plates were frozen at -80°C until citrate synthase activity was ready to be measured. Briefly, 113 μl of assay buffer (200 mM Tris buffer pH8.0 (First Base, BUF-1415-1L-pH8.0) supplemented with 0.2% Triton^TM^ X-100 (v/v) (Sigma-Aldrich, T8787), 100 μM DTNB (Sigma-Aldrich, D8130-5G) in 100 mM Tris buffer pH8.0, and 1 mM Acetyl-CoA (Santa Cruz Biotechnology, sc-214465C) was added to each well before 5 μl of 10 mM oxaloacetic acid (Sigma-Aldrich, O4126-1G) was assed as the reaction substrate. Absorbance at 412 nm was measured using a M200 Microplate Reader (Tecan) at 37 °C for a minimum time interval of 5 mins. Citrate synthase activity was subsequently calculated using the formula below.

Citrate synthase activity=Σ*i*=1*n*[(*Ai*−*A*0*ti*−*t*0)]/*n*⋅(*n*∈*N*∗)

where n is the total number of absorbance records, A is the absorbance recorded, and t is the time.

### Electron Transfer Chain Enzymatic Activity Assay

CI and CIII enzymatic activity in cultured BMDMs were determined by adapting reference protocols X where respective complex-specific substrates and electron acceptors were added to induce a colorimetric change. All measurements were recording using a M200 Microplate Reader (Tecan) at 37 °C with a minimum time interval for 20 mins. Both CI and CIII enzymatic activity were defined by the slope of their respective absorbance measurements against time.

### Mitochondrial Membrane Potential

Mitochondrial membrane potential was measured using Tetramethylrhodamine Ethyl Ester (TMRE) (Sigma-Aldrich, 87917). BMDMs were seeded onto 48-well plates at a density of 250,000 cells per well one day prior to the assay. 30 mins before the appropriate timepoints, TMRE was added to a final well concentration of 100 nM and allowed to incubate. Cells were then washed with warm PBS (Cytiva, SH30028.02), trypsinised, and pelleted before being resuspended in warm media for acquisition by flow cytometry.

### Overall Mitochondrial ROS Quantification

Total mitochondrial ROS was measured using MitoSOX^TM^ Mitochondrial Superoxide Indicators (Invitrogen, M36008). BMDMs were seeded onto 48-well plates at a density of 250,000 cells per well one day prior to the assay. 30 mins before the appropriate timepoints, MitoSOX^TM^ was added to a final well concentration of 5 μM and allowed to incubate. Cells were then washed with warm PBS, trypsinised, and pelleted before being resuspended in warm media for acquisition by flow cytometry.

### Site Specific ROS Quantification

ROS generation at specific sites within CI and CIII was determined by adapting reference protocols X. BMDMs were seeded onto 96-well black polystyrene microplates at a density of 80,000 cells per well one day prior to the assay. At the appropriate timepoint after infection or treatment, culture media was replaced with assay buffer containing 120 mM KCl (Sigma-Aldrich, P5405), 5 mM HEPES (Gibco, 15630080), 1 mM Ethylene glycol-bis(2-aminoethylether)-N,N,N′,N′-tetraacetic acid (EGTA) (Sigma-Aldrich, E4378), 12.5 U/mL superoxide dismutase (Scientific Laboratory Supplies, S7446-15KU), 2.5 U/mL horse raddish peroxidase (HRP) (Sigma-Aldrich, P8375), 12.5 µM Amplex UltraRed (Invitr), 0.3% w/v fatty acid-free free bovine serum albumin (BSA) (Sigma-Aldrich, A7030), and 0.0025% digitonin (Santa Cruz Biotechnology, sc-280675D). Superoxide formation at specific ROS sites within the ETC was stimulated with the use of complex-specific substrates and inhibitors. CI Q_R_ ROS was induced using 5 mM sodium succinate (Sigma-Aldrich, S7501) and 0.3 μM S3QEL 2 (Sigma-Aldrich, SML1554) but inhibited with 1 µM nigericin (MedChem Express, HY-100381). CIII Q_O_ ROS was induced using 5 mM sodium succinate, 2.5 μM antimycin and 4 μM rotenone but inhibited with 2 μM myxothiazol (Sigma-Aldrich, T5580). Fluorescence intensity (peak excitation/emission = 540 nm/590 nm) overtime was detected using an Infinite Lumi Microplate Reader (Tecan). ROS production was defined by the gradient of the slope after subtracting the corresponding measurement in the presence of the inhibitor. All measurements were normalised to citrate synthase activity.

### Inhibition of Site Specific ROS Generation

ROS generation at CI and CIII was pharmacologically inhibited with the use of S1QEL 1.1 (Sigma-Aldrich, SML1948) and S3QEL 2, respectively. Drugs were prepared in dimethyl sulfoxide (DMSO) and given to the cells at indicated concentrations 2 hours before infection as a pre-treatment. Fresh drug was given 2 hours after infection when the virus was washed off and media was replaced. Cells were then returned to the incubator until the time of harvest or assay. Untreated control cells were given DMSO.

### Mitochondria Isolation

Mitochondria was isolated from cultured BMDMs using reference protocols. Briefly, BMDMs from confluent 10 cm dishes were harvested by scraping, washed once with PBS, and pelleted. Cells were resuspended in mitochondria isolation Buffer A (83 mM sucrose (Sigma-Aldrich, S0389), 10mM HEPES, pH7.4) supplemented with 1x cOmplete EDTA-free protease inhibitor cocktail (Roche, 5056489001) and 1 mM phenylmethylsulfonyl fluoride (PMSF) (Sigma-Aldrich, 10837091001) for 10 min on ice before being homogenised using Potter-Elvehjem tissue grinders with PTFE pestles powered by a motorized drill for a total of 40 strokes. Efficiency of cell membrane lysis was confirmed using trypan blue under white field microscopy. Cell homogenates were spun at 1,000 x g for 10 mins at 4 °C and supernatants was collected in a new tube. The remaining cell pellets were resuspended in mitochondria isolation. Buffer B (250 mM sucrose, 10 mM HEPES pH7.4) supplemented with 1x cOmplete protease inhibitors and 1 mM PMSF for the extraction to be repeated. Supernatants from both extractions were combined and centrifuged at 12,000 x g for 15 mins at 4 °C. At the end, supernatants were aspirated to yield the mitochondria pellet, which was resuspended in mitochondria isolation Buffer C (320 mM sucrose, 10 mM Tris-HCL, 1 mM EDTA, pH7.4) supplemented with 1x cOmplete protease inhibitors and 1 mM PMSF for protein quantification using the Pierce^TM^ BCA kit (Thermo Scientific, 23225).

### Immunoblotting

30 μg of isolated mitochondria were lysed in RIPA buffer supplemented with 1x cOmplete EDTA-free protease inhibitor cocktail for 10 mins on ice. Any insoluble debris was removed by centrifugation at 20,000 x g for 10 mins. Thereafter, lysates were heated in 1X laemmli sample buffer (50 mM Tris-HCl pH 6.8, 2% SDS, 10% glycerol, 12.5 mM EDTA, 0.02% bromophenol blue, 50 mM DTT) for 5 mins at 95 °C. Proteins were resolved in Bolt™ Bis-Tris Plus 4-12 % gradient gels (ThermoFisher, NW04122BOX) and transferred onto 0.2 μM PVDF membranes (Bio-Rad, 1704272) using the semi-dry Trans-Blot Turbo Transfer System (Bio-Rad, 1704150). Membranes were subsequently blocked with 5% non-fat milk (Bio-Rad, 170-6404) in Tris Buffered Saline (TBS) (First Base, BUF-3030-20X4L)-Tween 20 0.1% (v/v) (Sigma-Aldrich, P1379) for 1 hr at room temperature before being immunoblotted with primary antibodies against proteins of interest overnight at 4 °C. The antibodies and the appropriate dilution factor used are listed in Table 2. Membranes were washed extensively in TBST and probed with HRP-conjugated secondary antibodies against either mouse (Invitrogen, 31430) or rabbit IgG (Jackson ImmunoResearch, 111-035-144) at 1:10,000 dilution and washed again. Chemiluminescence was captured by the Chemidoc MP imaging system (Bio-Rad, 12003154).

**Table 1:**
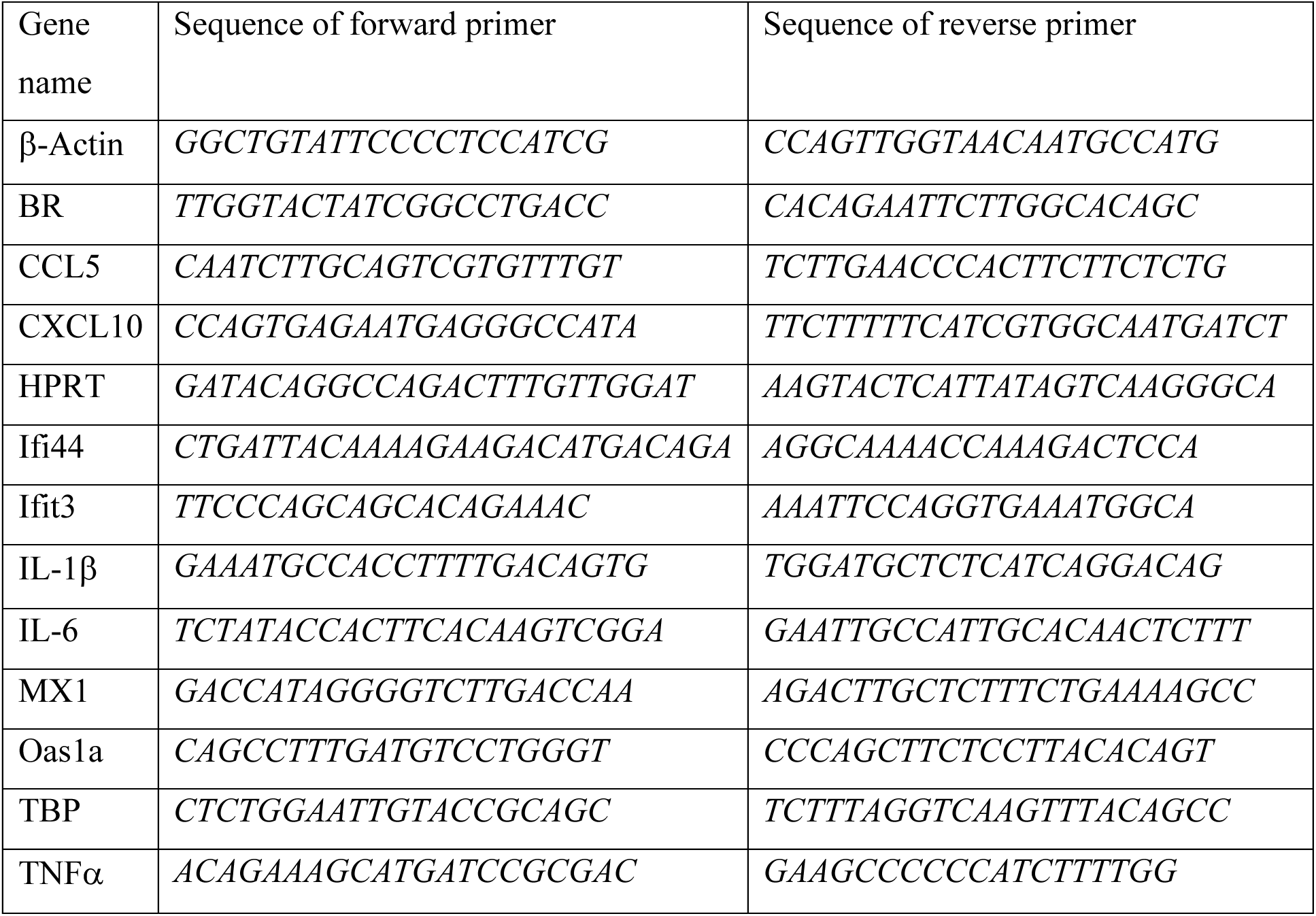
List of primer sequences used for qPCR.

**Table 2:** List of primary antibodies used for immunoblotting.

| Antibody | Dilution | Source | Identifier |
| --- | --- | --- | --- |
| Anti-β-Actin | 1:5,000 | Sigma-Aldrich | A2228 |
| Anti-BR | 1:2,000 | GL-Biochem (custom) |  |
| Anti-GAPDH | 1:5,000 | Cell Signalling Technology | 2118S |
| Anti-MT-CYB | 1:1,000 | Proteintech | 55090_I-AP |
| Anti-NDUFA9 | 1:1,000 | Thermo Fisher | 459100 |
| Anti-SDHA | 1:2,000 | ABclonal | A2594 |
| Anti-TOM70 | 1:2,000 | ABclonal | A21210 |
| Anti-UQCC1 | 1:2,000 | Sigma-Aldrich | HPA034875 |
| Anti-UQCC2 | 1:1,000 | ABclonal | A19955 |
| Anti-UQCRC1 | 1:3,000 | ABclonal | A3339 |
| Anti-UQCRFS1 | 1:2,000 | Abcam | Ab14746 |

### Blue-native Page

50 μg of isolated mitochondria were solubilised for 20 mins on ice in 1X NativePAGE^TM^ Sample Buffer (Invitrogen, BN2003) containing digitonin at a digitonin/protein ratio of 8 g/g. The insoluble fraction was removed by centrifugation at 20,000 x g for 20 mins. NativePAGE^TM^ 5% G-250 sample additive (Invitrogen, BN2004) was subsequently added to the solubilised protein lysate to a final concentration of 0.5% before loading into NativePAGE^TM^ 3-12% Bis-Tris protein gels (Invitrogen, BN1001BOX). Protein complexes were resolved by electrophoresis firstly for 30 mins at 150 V in dark blue cathode buffer and subsequently for 60 mins at 250 V in light blue cathode buffer. Dark blue cathode buffer was prepared by diluting 20X NativePAGE^TM^ Cathode Buffer Additive (Invitrogen, BN2002) to the appropriate working concentration with 1X NativePAGE^TM^ Running buffer (Invitrogen, BN2001). Light blue cathode buffer was derived by further diluting the dark blue cathode buffer 10 times with 1X NativePAGE^TM^ Running buffer. The separated protein complexes were transferred onto 0.2 μM PVDF membranes using the semi-dry Trans-Blot Turbo Transfer System. Membranes was subsequently fixed in 8% acetic acid for 10 mins, destained in methanol for 10 mins, and blocked with 5% non-fat milk in TBS-Tween 20 0.1% (v/v) for 1 hr at room temperature before being immunoblotted with primary antibodies against proteins of interest as described in the immunonblotting section above. The antibodies and the appropriate dilution factor used are listed in Table 2.

### Mass Spectrometry

100 μg of isolated mitochondria were resolubilised and denatured in 8 M urea prepared in 50mM Tris-HCl (pH8.5). Following sonication with a probe sonicator on ice, proteins were precipitated in acetone overnight at -20 °C and subsequently pelleted at 20,000 x g, and dried. Samples were next reduced with 10 mM tris(2-carboxyethyl)phosphine (TCEP) (GoldBiotechnology, TCEP) for 20 mins at 55 °C and alkylated with 55 mM chloroacetamide (Sigma-Aldrich, C0267) for 30 mins at 25 °C in the dark. Protein digestion was performed first using Lysyl Endopeptidase (Novachem, 129-02541) at an enzyme/protein ratio of 1:40 for 3 hr at 37 °C followed by digestion with mass spectrometry-grade trypsin protease (Thermo Scientific, 90058) at an enzyme/protein ratio of 1:20 overnight at 25 °C. Samples were then desalted using 1cc columns before being passed through the mass spectrometer.

### Mouse Husbandry

BR KO mouse lines were generated as previously described (Liang et al., 2022). All mice were housed in individually ventilated cages on a 12 hr light/dark cycle with water and normal chow diet provided ad libitum. Room temperature and relative humidity were maintained at 21-24 °C and 40-60%, respectively. 8- to 12-week-old mice were used in all experiments. All animal procedures were conducted in accordance with protocols approved by Duke-NUS, SingHealth Institutional Animal Care and Use Committee.

### Mouse Influenza Infection

Mouse-adapted influenza H1N1 virus A/PR/8/34 (PR8) was obtained from AVS Bio and the use of the virus in vivo was approved by and carried out in compliance with RCULAC and IACUC regulations (Singhealth). Mice were anaesthetised and inoculated intranasally with 40 μl of virus diluted in PBS. Sublethal infection was achieved using 250 PFU/mouse whilst lethal/severe infection was achieved using 1250 PFU/mouse. Control mice received 40 μl of sterile PBS intranasally. Mice were weighed and observed for clinical signs of disease daily for 21 days or until sacrifice at the respective timepoints.

### Viral Titer Quantification

Lung viral load was determined using probe-specific qPCR of total RNA isolated from lung homogenates. qPCR was performed using Luna Probe One-Step RT-qPCR Mix (New England Biolabs, E3005X) with PR8 polymerase (PA) gene-specific primers and probe (Integrated DNA Technologies) in a CFX384 Touch Real-Time PCR Detection System (Bio-Rad Laboratories, 1855484). The forward and reverse primers are *CGGTCCAAATTCCTGCTGA* **and** *CATTGGGTTCCTTCCATCCA*, respectively and the probe sequence is *CCAAGTCATGAAGGAGAGGGAATACCGCT*.

### Cytokine Analysis

Virus-stimulated cytokine production in BMDMs and mouse lung was analysed using the LEGENDplex^TM^ Multianalyte Immunoassay (BioLegend, Cat. No. 740621) according to the manufacturer’s instructions. For in vitro quantification, cells were seeded in 24-well plates at a density of 250,000 cells/well. Supernatants were subsequently harvested at respective timepoints and centrifuged at 1,000 x g for 5 mins to remove debris for use in the assay. For in vivo quantification, lung homogenates were assayed and prepared by homogenising whole lung tissue in homogenisation buffer (0.5% Triton X-100, 150mM NaCl (Sigma-Aldrich, S9625-5KG), 15mM Tris, and 2.5mM MgCl_2_ (Sigma-Aldrich, M2670-100G) supplemented with 1x cOmplete protease inhibitors and 1mM PMSF at pH7.4) using a microtube homogeniser. Crude lung homogenates were centrifuged at 10,000 x g for 10 mins at 4 °C to obtain cell-free supernatants that were used in the assay. The levels of cytokines were analysed using the LEGENDplex^TM^ v8.0 software and normalised to RNA yield.

### Histology

The post caval lobe of the lung was dissected and fixed in 10% neutral buffered formalin solution (Sigma Aldrich, HT501128) at 4 °C overnight. Following which, samples were dehydrated in 70% ethyl ethanol before being embedded in paraffin wax. 5 μm tissue sections were obtained and stained with haematoxylin and eosin. All images were taken by an Olympus BX53 light microscope at 10 x magnification.

### Lung Dissociation

PR8-infected and control mice were sacrificed by CO_2_ asphyxiation and their lungs were perfused with PBS through the right ventricle before removal. Lung tissues were crudely minced and resuspended in 2 ml of digestion buffer (RPMI (HyClone, SH30027.01) containing 0.4 mg/ml Liberase DL Research Grade (Roche, 5466202001), 20 U DNase I (Roche, 4716728001), and 1% fatty acid-free BSA before being transferred into gentleMACS^TM^ C Tubes (Miltenyi Biotech, 130-093-237). Crude lung tissues were further dissociated by running the *lung1* and *lung2* programs on the gentleMACS Octo Dissociator (Miltenyi Biotech, 130-096-427) with a 30 min digestion at 37 °C in between. Digestion was quenched with PBS supplemented with 5% FBS and 1mM EDTA. The samples were subsequently passed through 70 μm strainers (SPL Life Sciences, 93070) and pelleted by centrifuge at 500 x g for 10 mins for erythrocyte lysis using a commercial lysis buffer (BioLegend, 420301). Cells were lastly resuspended in FACS buffer (PBS supplemented with 2% FBS and 1mm EDTA) for either flow cytometry or the single cell RNAseq workflow.

### Flow Cytometry

Single cell suspensions for flow cytometry were obtained from mouse lung as described above. Cells were pelleted at 500 x g for 5 mins and first treated with FC block (BD Biosciences, 553142) at 1:100 dilution for 15 mins at 4 °C. Cells were washed once with FACS buffer (PBS supplemented with 2% FBS and 1 mm EDTA) before incubation with a antibody cocktail containing LIVE/DEAD 405 (Thermofisher), CD45.2, CD8, CD11b, CD11c, Ly6c, Ly6g (BioLegend) at 1:200 dilution for 30 mins at 4 °C. Following three washes with FACS buffer, cells were fixed in 4% paraformaldehyde (w/v) (Thermo Scientific, 28906) for 15 mins at 4 °C. Lastly, cells were resuspended in FACS buffer for acquisition by LSRFortessa^TM^ Cell Analyser (BD Biosciences, 23-11617). Data acquired was analysed using FlowJo version 10.7.1. Gating strategy for lung monocytes are is as follows:

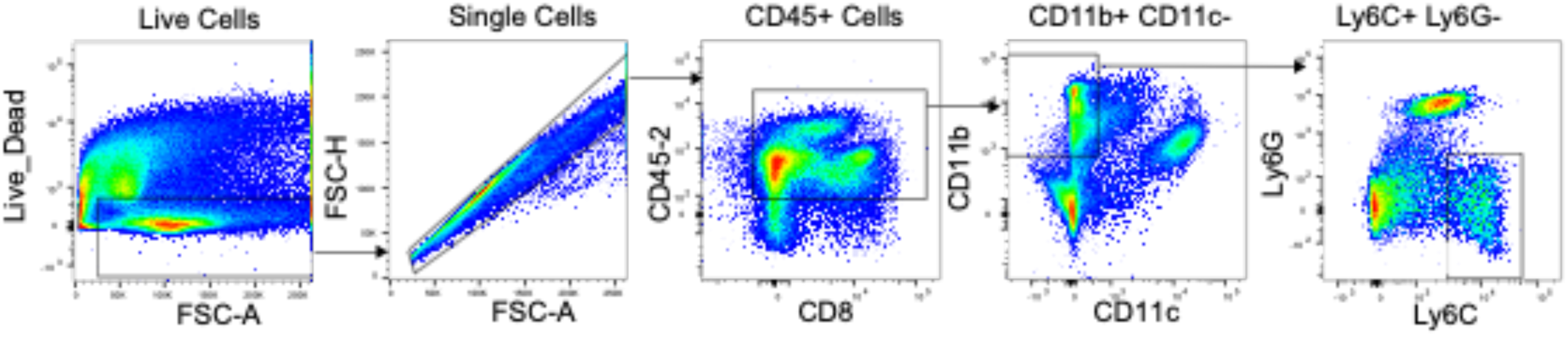

### Preparation of Single Cell RNAseq Libraries

Single cell suspensions obtained from lung dissociation as described in the lung dissociation section above were further enriched for viable cells by dead cell removal using a commercial dead cell removal kit (Miltenyi Biotech, 130-090-101) and collecting the flow through from MS columns (Miltenyi Biotech, 130-042-201). The flow through was subsequently strained through 40 μm Flowmi® Cell Strainers (Bel-Art™, BAH136800040) before proceeding with the Chromium Next GEM Single Cell 3’ Reagent kit v3.1 (10x Genomics, 1000269) workflow according to the manufacturer’s instructions.

### Single Cell RNAseq Analysis

The single cell libraries were first mapped with Cellranger v6.1.1 (Zheng et al., 2017) to the mouse genome(mm10) and then dehashed with HtoDemux function in Seurat 4.0 (Hao et al., 2021). Cells that have less than 300 or more than 6500 unique features are filtered as low-quality cells. Cells with mitochondrial counts higher than 15 percent are also filtered. The data was then normalized and scaled with SCTransform (Hafemeister & Satija, 2019) where 3000 highly variable genes were chosen. The different datasets were then batched corrected with Seurat Integration (Stuart et al., 2019) and dimension reduction is conducted with principal component analysis (PCA). We ran UMAP and Louvain (McInnes et al., 2018)clustering on 20 PCs based on the cutoff from the elbow plot. Differential expression was conducted using the Wilcoxon rank test to identify markers genes which were used to annotate the clusters.

To understand the change in cellular states across different timepoint and different conditions, we make use of edgeR (Robinson et al., 2010)to identify the differentially expressed genes between both different timepoint and conditions. Gene set enrichment analysis (GSEA) was conducted on the differentially expressed genes using the fgsea (Korotkevich et al., 2019) package along with the MsigDB database to identify associated biological pathways and processes.

To study the changes in composition between the different Br WT and KO, we made use of MILO, a statistical framework which detects compositional changes at neighborhood level. We provide MILO with the PCA embedding and the condition labels and ran the downstream with default parameters except for specifying the number of neighbors to be 25 when building the neighborhood graph.

Viral-Track (Bost et al., 2020), a computational method designed for identifying viral presence in single-cell data by scanning the unmapped scRNA-seq data for viral RNA is used for identification of the Pr8 Virus. We used the Pr8 virus obtained from viruSITE (Stano et al., 2016) as the reference genome.

We then performed cell-to-cell interaction analysis with CellChat (Jin et al., 2021)which made use of celltype annotations and counts data. The default database for mouse is used. We ran CellChat on each of the conditions separately before merging them for further downstream plotting. Lastly, we extracted out the classical monocytes and inflammatory monocytes from our single cell data to run Compass which allows us to study the metabolic difference between Br WT and KO. We ran compass with the default parameters. We then use Cohen’s d on the output metabolic reaction scores to quantify the effect size between the two conditions. Gene module scores were calculated using the AddModuleScore function in Seurat (package version 5.3.0). M1-like inflammatory module and fatty acid oxidization (FAO) module were defined using the following canonical gene sets respectively: *Il1b, Il6, Tnf, Cxcl9, Cxcl10, Nos2, Irf5, Stat1, Cd80, Cd86* and *Cpt1a, Acadvl, Acadm, Ppargc1a*. For pseudotime analysis of inflammatory monocyte trajectories, classical and inflammatory classical monocytes were subset and analyzed using Monocle 3 (package version 1.3.7). An inflammatory gene module (*Ly6c2, Fcgr1, Isg15*) was computed. Cells were normalized using SCTransform, followed by dimensionality reduction, clustering, and UMAP embedding in Seurat. Trajectory learning was performed using Monocle 3’s graph-based approach, and pseudotime was inferred by ordering cells from a low-inflammatory root cluster based on inflammatory module score. Pseudotime values for each cell were mapped back onto the Seurat object for visualization between WT and Br KO. *Monocyte isolation*

Mouse monocytes were isolated and derived using reference protocols. Briefly, bone marrow cells were harvested by flushing sterilised femurs and tibias of BR WT and KO C57BL/6 mice, followed by erythrocyte lysis using a commercial lysis buffer. Primary monocytes were subsequently obtained by negative selection using the EasySep™ Mouse Monocyte Isolation Kit (STEMCELL Technologies, 19861) and running the cell suspension through MS columns. Isolated monocytes were maintained in suspension in DMEM supplemented with 10% FBS, 1% penicillin-streptomycin, and 25ng/ml granulocyte-macrophage colony stimulating factor (GM-CSF) (Sigma-Aldrich, SRP3201).

### Transwell Migration Assay

Monocyte migration was analysed by a transwell migration assay using 5 μm pore size polycarbonate membrane Transwell tissue culture inserts (Corning, CLS3421-48EA). MLE-12 cells or BMDMs were seeded in the chamber at a density of 550,000 cells/well and 250,000 cells/well, respectively. On the day of the assay, chamber cells were infected with the PR8 virus at an MOI of 10. Monocytes in DMEM supplemented 10% FBS, 1% P/S, and 25ng/ml GM-CSF were subsequently seeded into the transwell inserts at a density of 180,000 cells/insert. Transwell inserts were transferred into the respective chambers 8 hr after the start of infection and returned to a 37 °C 5% carbon dioxide incubator. Monocytes were allowed to migrate for 5 hours, after which cells were fixed for quantification. Non-adherent monocytes that had migrated into the chamber were collected by harvesting the media in the lower chamber, staining for Ly6C, and acquiring the cells by flow cytometry using the LSRFortessa^TM^ Cell Analyser. Cell number was calculated and normalised based on spike-in beads. Adherent monocytes that had migrated but remained attached to the bottom side of the transwell were fixed with 4% paraformaldehyde (w/v) and stained with Hoescht at 1:1000 dilution. Cells on the transwell insert that had not migrated were rubbed off with a wet cotton tip. The remaining cells were visualised using a Leica DMi8 fluorescent microscope at 20 x magnification and 4 non-overlapping representative images were taken per well.

